# Reconstructing and comparing signal transduction networks from single cell protein quantification data

**DOI:** 10.1101/2024.03.29.587331

**Authors:** Tim Stohn, Roderick van Eijl, Klaas W. Mulder, Lodewyk F.A. Wessels, Evert Bosdriesz

## Abstract

**Motivation:** Signal transduction networks regulate a multitude of essential biological processes and are frequently aberrated in diseases such as cancer. Developing a mechanistic understanding of such networks is essential to understand disease or cell population specific signaling and to design effective treatment strategies. Typically, such networks are computationally reconstructed based on systematic perturbation experiments, followed by quantification of signaling protein activity. Recent technological advances now allow for the quantification of the activity of many (signaling) proteins simultaneously in single cells. This makes it feasible to reconstruct signaling networks from single cell data.

**Results:** Here we introduce single cell Comparative Network Reconstruction (scCNR) to derive signal transduction networks by exploiting the heterogeneity of single cell (phospho)protein measurements. scCNR treats stochastic variation in total protein abundances as natural perturbation experiments, whose effects propagate through the network. scCNR reconstructs cell population specific networks of the same underlying topology for cells from diverse populations. We extensively validated scCNR on simulated single cell data, and we applied it to a dataset of EGFR-inhibitor treated keratinocytes to recover signaling differences downstream of EGFR and in protein interactions associated with proliferation. scCNR will help to unravel the mechanistic signaling differences between cell populations by making use of single-cell data, and will subsequently guide the development of well-informed treatment strategies.

**Availability and implementation:** scCNR is available as a python module at https://github.com/ibivu/scmra. Additionally, code to reproduce all figures is available at https://github.com/tstohn/scmra_analysis.

**Supplementary information:** Supplementary information and data are available at Bioinformatics online.

## 1 Introduction

Signal transduction networks play a vital role in cell physiology, where they regulate various biological processes like differentiation, proliferation, and apoptosis. Extracellular signals are commonly transmitted through intracellular networks of interacting proteins called kinases, which activate each other through post-translational modifications such as phosphorylation. Diseases such as cancer often are a consequence of aberrations in this signaling machinery (Hanahan and Weinberg, 2011; Kolch *et al*., 2015), and many cancer drugs specifically target proteins within the signaling networks. However, adaptation and rewiring of the network (Ahronian *et al*., 2015) in response to treatment often limits the durability of a clinical response (Gerosa *et al*., 2020; Hoffman *et al*., 2023; Tognetti *et al*., 2021). Obtaining a mechanistic (Does protein A activate protein B?) and quantitative (How strong is the influence of protein A on protein B?) understanding of those networks, and how they differ between cell populations (such as resistant, mutant cells), is a key challenge in cellular biology and has important implications for the design of treatment strategies (Bosdriesz *et al*., 2022; Klinger *et al*., 2013).

Various methods have been developed to solve this problem. Ordinary differential equations based models are detailed and quantitative (Fey *et al*., 2015; Raue *et al*., 2015), but rely on numerous measurements and are restricted to small systems. Boolean logical network models have simpler formulations, but can only represent non-quantitative network interactions and are often not able to explain cyclic structures and feedback-loops (Oates *et al*., 2012; Grieco *et al*., 2013; Saez-Rodriguez *et al*., 2009; Hill *et al*., 2012). Modular response analysis (MRA) finds the middle ground by modelling networks quantitatively without the complexity of fully dynamical models (Kholodenko *et al*., 2002; Bruggeman *et al*., 2002). MRA determines the interaction strengths between proteins based on systematic perturbation experiments, in which the states of all nodes in the network are recorded before and after the perturbations. MRA is able to detect cross-talks and feedback loops, and has been successfully employed to quantify novel network topologies (Dorel *et al*., 2018; Klinger *et al*., 2013). Various alternative formulations have been developed, such as optimisation based approaches (Bosdriesz *et al*., 2018), maximum-likelihood approaches (Klinger *et al*., 2013; Ahlmann-Eltze and Huber, 2023) or Bayesian methods (Halasz *et al*., 2016; Santra *et al*., 2013; Rukhlenko *et al*., 2022). While most methods model signaling networks in a specific context, differences between networks derived from different contexts are often most informative. For example, comparing networks derived from a cell line before and after acquiring resistance to a targeted inhibitor can aid in elucidating the signaling changes that drive the resistance mechanism (Bosdriesz *et al*., 2018), models of cell lines with and without a particular oncogenic mutation can help in prioritizing drug combinations that are specific to a particular genetic background (Bosdriesz *et al*., 2022), and modeling the dependence of a signaling network on cell-states can help predict interventions that control cell fate decisions (Rukhlenko *et al*., 2022). To compare networks across contexts, we recently developed Comparative Network Reconstruction (CNR) (Bosdriesz *et al*., 2018), which aims to identify quantitative differences between the signaling networks derived from cell populations in different contexts.

Most methods that model signal transduction networks were developed for bulk data. However, even in isogenic populations in a homogeneous environment, cells in different cell states are known to respond differently to the same instructions (Kim *et al*., 2018; Wang *et al*., 2022; Kramer *et al*., 2022). This has important implications for drug resistance and cancer treatment design (Korkut *et al*., 2015; Aissa *et al*., 2021) and efforts have been made to link signaling networks to their effect on the cell state transition in perturbed RAF inhibitor-resistant cancer cells (Rukhlenko *et al*., 2022). Obtaining such insights has long been elusive because we lacked the right data to study them, but recent development of technologies for high dimensional single-cell quantification of (phospho-)proteins and post-translational modifications, based on mass cytometry (Tracey *et al*., 2021), DNA-barcoded antibodies (Eijl *et al*., 2018; Stoeckius *et al*., 2017; Sheng *et al*., 2022) and spatial methods such as iterative indirect immunofluorescence imaging (Gut *et al*., 2018) now allow us for the first time to study and model the mechanisms underlying the heterogeneity of signal transduction in a data-driven manner.

Here, we describe single-cell Modular Response Analysis (scMRA) and single-cell Comparative Network Reconstruction (scCNR). scMRA is a method to infer signaling networks from single-cell quantification of active and total protein abundance. Importantly, it exploits stochastic variability of total protein counts between cells as ‘natural perturbation experiments’, similar to perturbations at the bulk level in classical MRA (Kholodenko *et al*., 2002), thereby eliminating the need for extensive perturbation experiments. By considering the data captured from each single cell as a data-point, scMRA vastly increases the number of observations from which the reconstructions are derived. Typically, the signaling *diversity* between cell populations is of most interest, e.g. due to cell state effects such as the cell cycle and differentiation, or due to treatment effects or disease progression or the emergence of resistant sub-populations. Therefore, we extended scMRA to single-cell Comparative Network Reconstruction (scCNR), in order to identify which interactions differ quantitatively between cell populations. Similar to CNR (Bosdriesz *et al*., 2018), scCNR reconstructs a single shared network topology with cell population-specific interaction strengths. We extensively validate scMRA and scCNR on simulated single cell data of the epidermal growth factor receptor (EGFR) signaling pathway and showed that both methods perform well using as few as hundred cells as input, and in the presence of considerable noise. Furthermore, we applied scCNR to a dataset where we quantified 70 (phospho)proteins of key signaling nodes using single-cell ID-seq (Eijl *et al*., 2018) in EGFR-inhibitor treated keratinocytes, and showed that scCNR recovers meaningful biological diverged protein interactions downstream of EGFR and related to proliferation.

## 2 Materials and Methods

### 2.1. Formulation of single-cell MRA and CNR

Here, we briefly describe the formalism underlying single-cell Modular Response Analysis (scMRA) and single-cell Comparative Network Reconstruction (scCNR). More details and a full derivation of the equations can be found in the Supplementary Information 5.2.

scMRA and scCNR exploit stochastic variation in protein abundances to identify and quantify the interaction strengths between different nodes in a signal transduction network. As input, they require single-cell measurements of the abundance and activity of all nodes in the network, and potentially the applied perturbations. These are related through the following set of linear equations (one for each protein *i* in each cell *a*):

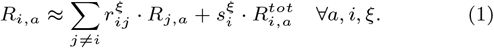

where 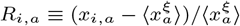 is the deviation of the measured abundance of active protein *x* from the population average, 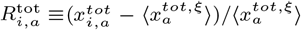 is the deviation of the measured total protein abundance *x*^*tot*^ from the population average, 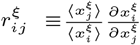 is the population-specific interaction strength between node *j* and node *i*, with *r*_*ii*_ *≡ −*1, s the population-specific sensitivity of node *i* activity to a change in total protein, and 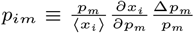 is the direct effect that perturbation *m* has on the activity of node *i*.

scMRA and scCNR both solve an MIQP optimization problem that aims find the values of 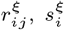 and *p*_*im*_ that minimize the squared error *ϵ* between the measured active protein abundance per cell and the prediction of the model as described by equation (1). Additionally, two *L*_0_-regularization penalties are added: one for the number of edges in the network, and one for the number of population-specific interaction strengths, sensitivities to change in total protein and perturbation effects. The full MIQP formulation then reads as follows:

minimize:

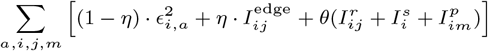

Subject to :

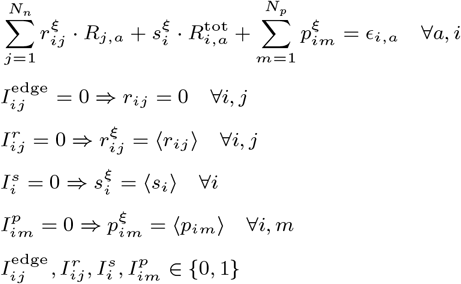

where 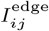 is a binary indicator for the presence or absence of an edge between nodes *i* and *j*. 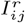 and 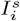 are binary indicators for a population-specific interaction strength and sensitivity to total protein changes, respectively, and are only present for scCNR. The hyperparameter *η* and *θ* determine the relative weight of the regularization penalties. For the inclusion of perturbation effects see supplementary 5.4. The optimization problem was solved using IBM ILOG CPLEX solver (version 20.1.0). CPLEX is available free of charge for academic purposes and guarantees an optimal solution within small numerical tolerances for our problem.

### 2.2. Simulating single-cell protein data

Single-cell protein abundance data was simulated using an existing ODE-based dynamic model of the EGFR signalling pathway (Orton *et al*., 2009). Single-cell total protein abundances were sampled from a log-normal distribution (*µ* = 0, *σ* = 0.1) and used to calculate the resulting quasi-steady state. Drug perturbations data was simulated as a 25% decrease in the catalytic activity of all protein-activating reactions. BRAF and RAS mutantions were modelled as a 100-fold reduction in the deactivation rate of active BRAF and RAF, respectively. Noise was added to the input data by multiplying the data with a random number drawn from a normal distribution with zero mean and a standard deviation equal to the noise. The ground-thruth interaction strengths between proteins of the Orton model were numerically calculated as the partial derivatives of active protein with respect to its upstream active protein.

### 2.3. scID-seq analysis

For full detail, see Supplementary Methods sections 5.1.2-5.1.6. Briefly, Primary pooled human epidermal stem cells where treated with the EGFR inhibitor AG1478 or vehicle (DMSO) for 48 hours. Subsequetly, the abundance of 70 phosphorylated and total proteins in 282 cells was quantified using single-cell ID-seq (Eijl *et al*., 2018). We ran scCNR to detect signaling differences between the untreated and EGFR-inhibitor treated cell population and used a prior literature derived network topology.

## 3 Results

### 3.1. scMRA reconstructs networks from total protein variation in single-cell protein data

Single cell Modular Response Analysis (scMRA) reconstructs signal transduction networks from the quantification of phospho- and total proteins in single cells. scMRA exploits the stochastic variability of total protein levels between single cells as natural perturbations to the signaling network (Fig. 1A). Stochastic differences in the abundance of total proteins between cells directly affect the abundance of the corresponding phosphoproteins. These changes in phosphoprotein levels then propagate through the network leading to distinct steady states of the network in each individual cell. Each cell is considered as an independent measurement of the underlying signaling network.

**Fig. 1:**
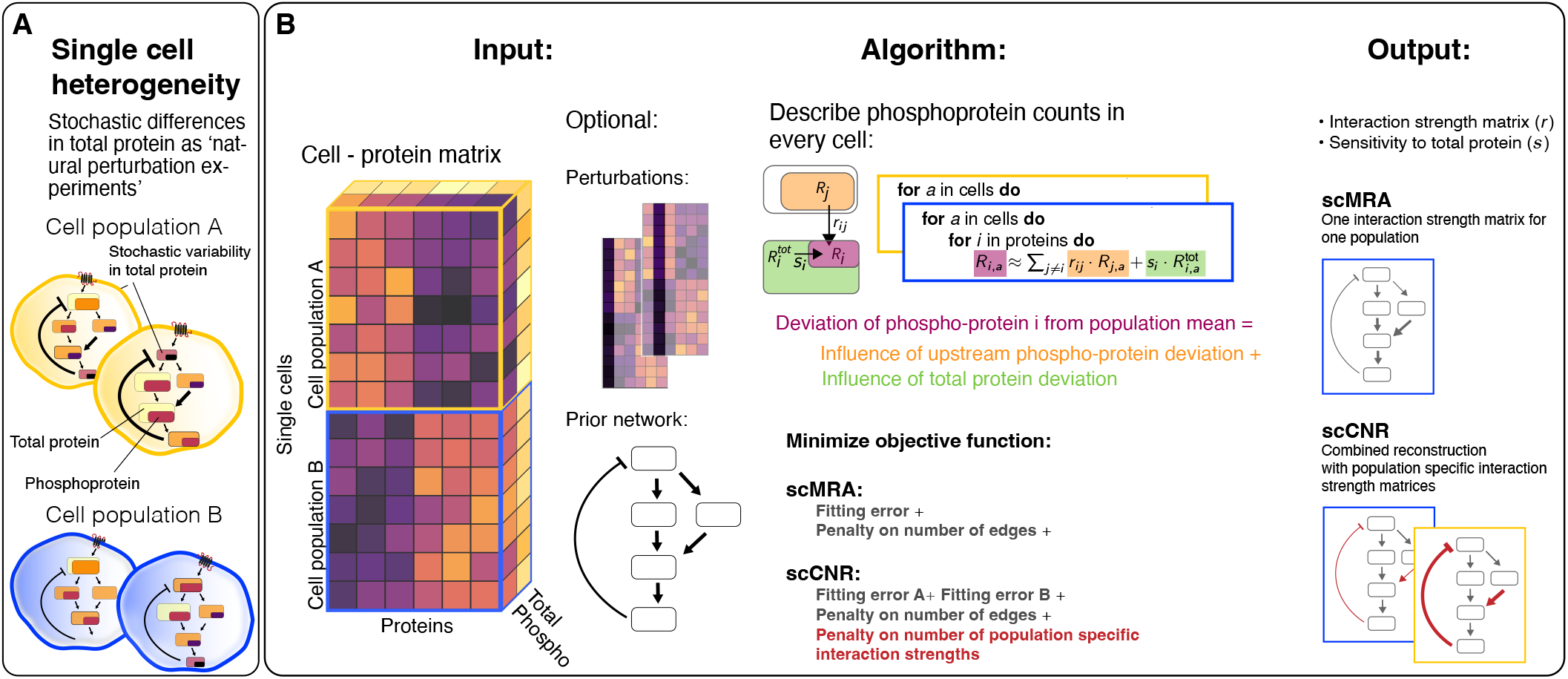
Schematic overview of scMRA and scCNR. (**A**) The methods exploit natural heterogeneity of phospho- and total protein abundances between cells to infer the network topology and quantify the interaction strengths between network nodes. (**B**) The methods take phospho- and total protein abundances from single cells as input, with additional cell population annotations for scCNR. Optionally the methods can be enriched with perturbation data and a prior network topology. The algorithms exploit deviations of total protein from the population mean (*R*^tot^) as ‘natural perturbations’. They fit the data to describe single-cell deviations of phosphoprotein from its population mean (*R*) for a single population (scMRA) or several populations in parallel (scCNR) to derive (cell population-specific) interaction strengths (*r*). The algorithms penalize the number of edges in the network. scCNR further penalizes the number of population-specific interaction strengths.

scMRA reconstructs a unique set of interaction strengths for a cell population (yellow and blue boxes representing populations A and B in Fig. 1). To model the variation between cell populations (which may represent for instance different cell states, cells with and without an oncogenic mutation, cells before and after acquiring resistance to a drug, or cells that are cultured for a long time in the presence or absence of an inhibitor), we developed single cell Comparative Network Reconstruction (scCNR) to identify population specific interaction strengths, just as CNR models condition-dependent interaction strengths in the bulk setting (Bosdriesz *et al*., 2018). This allows for the identification of the most relevant differences between the cell populations.

Input to the methods are deviations of abundances of total (*R*^tot^) and phosphoprotein (*R*) of each cell from the cell population mean, for each node in the network (Fig. 1B, ‘Input’). The output is the network topology described by interaction strengths between phosphoproteins (*r*), and sensitivities of phosphoproteins to deviations in its total protein (*s*) (Fig. 1B, ‘Output’). scMRA is formulated as a MIQP problem and fits a model that (i) for each node in each cell aims to explain the deviations of phosphoprotein abundance from the population mean and (ii) penalizes the model complexity (number of interactions) to derive a core signaling network. scCNR also derives a core signaling network with a small number of interaction strength differences between cell populations, while still producing a good model fit by (iii) penalizing the number of population-specific interaction strengths (Fig. 1B, ‘Algorithm’).

Often, not all total protein abundance measurements are available for all nodes in the network. However, additional perturbations, e.g. by small molecular inhibitors, can be included in the experimental design and model to facilitate network reconstruction. Furthermore, the formulation of the algorithm allows for easy integration of prior network information for cases where the topology might be established and the main interest lies in quantifying the interactions (Fig. 1B, ‘Input’).

### 3.2. scMRA faithfully reconstructs the topology of signal transduction networks

To evaluate how well scMRA recovers network topology and node interaction strengths, we first set out to test it on simulated data for which the ground truth is known. To this end, we simulated single-cell data for the epidermal growth factor receptor (EGFR) signaling pathway using a previously published dynamic model described by ordinary differential equations (Orton model), and reconstructed the network (Orton *et al*., 2009). Note that since scMRA and scCNR aim to explain the steady-state deviation of phosphoprotein abundance from the cell population mean only, the scMRA and scCNR network reconstructions will be much simpler than the original model. Nevertheless, the true interactions between active proteins are unambiguously defined (Fig. 2A). We simulated single cells by sampling total protein abundances from a log-normal distribution, and calculated the corresponding phosphoprotein abundances at the steady state.

**Fig. 2:**
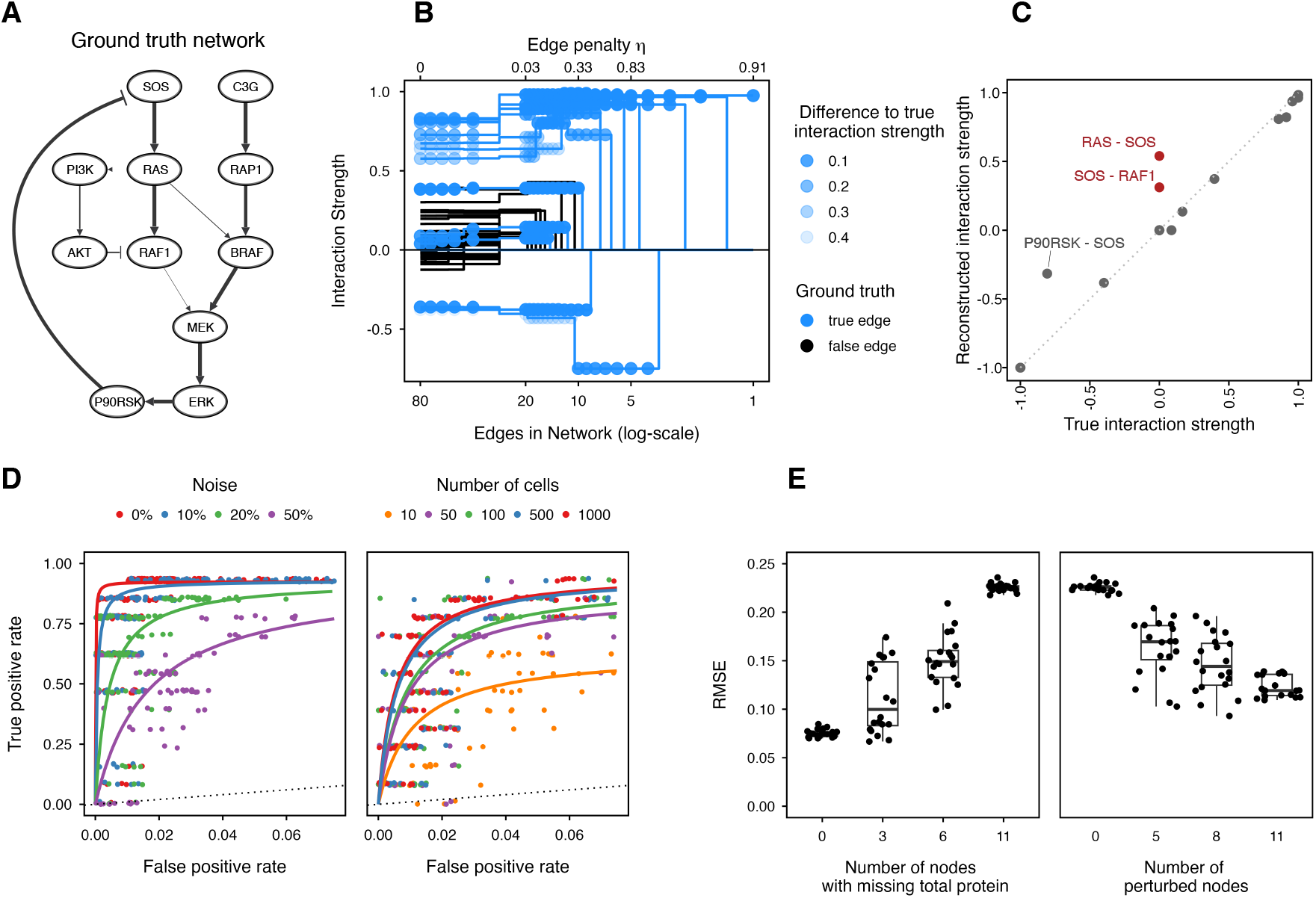
Evaluation of scMRA. (**A**) The EGFR signaling pathway according to the Orton Model. Arrow width is equivalent to the interaction strength derived from the Orton Model. (**B**) Interaction strengths reconstructed using scMRA for reconstructions of decreasing network complexity (increasing edge penalty *η*). True positives are indicated in blue and false positives in black. (**C**) Correlation between true and reconstructed interaction strengths for a network reconstruction with 1000 cells and 20% noise. (**D**) Receiver-operating characteristic curves for the effect of noise and number of cells on the reconstruction of the network topology. (**E**) RMSE between true and reconstructed interaction strengths for simulations with randomly removed total protein measurements and simulations with completely absent total protein measurements but additional perturbations for randomly selected nodes.

As a first test, we ran simulated 1000 cells, and added 20% noise to the data. We repeatedly reconstructed the signaling network using scMRA and progressively increased the complexity penalty *η*, resulting in increasingly sparse networks. Fig. 2B shows the reconstructed interaction strengths of the network, with true positive edges indicated in blue and false positives in black. Importantly, when increasing the *η*-penalty, false positive edges are the first to be eliminated, while the majority of true positive edges are retained. Furthermore, in reconstructions with low *η*-penalties and thus many edges, the false positive edges have typically weaker interaction strengths than the true positive edges. However, in reconstructions with a low *η*-penalty the strengths of the true positive edges are typically slightly different from their true values, as indicated by the alpha value of the points in Fig. 2B. As the penalty increases and false positive edges vanish, the reconstructed true edges in the network converge towards their actual interaction strength. This is further exemplified by the strong correlation between reconstructed and true interaction strengths (Fig. 2C).

### 3.3. scMRA works well in the presence of noise and with few input cells

As single-cell data is inherently noisy, we investigated how this impacts the performance of scMRA by adding varying amounts of noise on the input data. Similar to the analysis described above, we repeatedly reconstructed the Orton network while decreasing the network connectivity (by increasing the *η*parameter). Comparing the recovered edges to the ground truth network we generated receiver-operating characteristic (ROC) curves. With 20% noise a true positive rate of *>*75% can be achieved while keeping the false positive rate below 2.5% (Fig. 2D, left panel). Even with as much as 50% noise added to the data, a 70% true positive rate is attained at a false positive rate *<*6%. To put the numbers into perspective: a reconstructed network with 13 edges from a simulation with 1000 cells and 20% noise yields a network with two false positive edges. In addition to successfully recovering the network topology, the reconstructed interaction strengths are very similar to their true values, and only two edges with weak interaction strengths are not recovered (Fig. 2C).

Profiling large numbers of cells is not always feasible, as for instance even in large datasets specific cell populations of interest might be under-represented. Hence, we tested the influence of the number of cells measured on the performance of scMRA by simulating populations of various sizes, each with 20% noise added. The resulting ROC-curves are shown in Fig. 2D, right panel. For 500 cells or more, a true positive rate of *>*75% with a 2.5% false positive rate is attained. With as few as 50 cells a true positive rate of 75% with a 5% false positive rate can be attained. Taken together, this illustrates that scMRA can accurately reconstruct the network from few cells in the presence of considerable noise.

### 3.4. Perturbation experiments can compensate for missing total protein measurements

Ideally, scMRA uses measurements of phospho- and total protein for all nodes in the network, but this might not always be feasible due to experimental constraints. To assess the sensitivity of scMRA to missing total protein measurements we removed total protein information of randomly selected proteins from the simulated data. To quantify the reconstruction accuracy we calculated the root-mean-square error (RMSE) of the true and reconstructed interaction strengths, from simulations of 1000 cells with 20% noise added. To allow a fair comparison between reconstructed networks we compared networks with 13 edges.

As expected, increasing the number of proteins for which their total abundance is missing increases the RMSE of interaction strength reconstructions (Fig. 2E, left panel) from a mean RMSE of 0.07 when all total proteins are measured to 0.22 when they are all missing. To compensate for missing total protein measurements, scMRA can take perturbation data as input to improve network identification. To examine to which extent this could help to mitigate the negative effect of missing total protein, we simulated perturbations to a node as a 25% reduction in the catalytic activity of reactions activating that node. We randomly selected nodes to be perturbed, performed the perturbations and then reconstructed the network. When including the perturbations in the reconstruction, we kept the total number of cells to 1000. Adding perturbation data can decrease the RMSE from an average of 0.22 to 0.12 in the scenario with complete absence of total protein, but with perturbation data for all nodes in the network (Fig. 2E, right panel). This confirms that additional perturbations can partially compensate for missing total protein measurements and could be an option to include in the experimental design when total protein measurements might not be fully available.

### 3.5. scCNR identifies signaling differences between simulated wild-type and mutant cell populations

Tissue samples and cell cultures are inherently heterogeneous and can contain various cell types and states. To enable the investigation of differences in signaling between such populations we developed single cell Comparative Network Reconstruction (scCNR). scCNR is based on Comparative Network Reconstruction(Bosdriesz *et al*., 2018) that aims to identify what the most important quantitative differences are in signaling between cell populations. To this end, in scCNR we reconstruct a single network topology for all populations simultaneously, while allowing distinct interaction strengths for each population. scCNR outputs one shared network topology with a set of population-specific interaction strengths. In addition to the complexity penalty on the number of edges (*η*), scCNR also penalizes the number of population-specific interaction strengths (*θ*). This greatly reduces the total number of model parameters, thereby reducing overfitting and focusing on the most important differences. An additional advantage of this approach is that the reconstruction of each population is based on the full dataset, hence allowing information to be “borrowed” between cell populations.

We evaluated scCNR on simulated single cell populations of mutant RAS and BRAF cells, which we compared to the simulated wild-type population. These mutations cause constitutive activation of RAS and BRAF, respectively, which we modeled similarly to Orton et al. by a 100-fold reduction of the deactivation rate of the mutant protein (Orton *et al*., 2009). We simulated 250 cells per population and added 20% noise. Next, we reconstructed the signaling networks and compared them to the ground truth network of the Orton model. We estimated an appropriate value for *θ*, which reduces the mean squared error of the objective function while keeping the model complexity low (Fig. S2). This resulted in approximately four edges that differed between the mutant and wild-type populations.

The simulations of the wild-type and RAS mutant cell population result in major differences upstream of BRAF, which scCNR recovered truthfully (Fig. 3A and Fig. S3A). The BRAF mutant network, on the other hand, mainly differed downstream of BRAF within the BRAF-MEK-ERK-P90RSK cascade, which scCNR identified correctly as well (Fig. S4A and Fig. S3D). To quantitatively compare the scCNR reconstruction to the ground truth network, we reconstructed the signaling network for the wild-type and mutant cell population 20 times with 250 cells per population and 20% noise. Similar to previous simulations we aimed to reconstruct networks with 13 edges. The average reconstructed interaction strengths for both the RAS mutant and BRAF mutant cell populations correlated well with the true values for most edges (Fig. 3B and Fig. S4B). With the restriction on detecting four population-specific interaction strengths, small interaction strength differences (such as BRAF-MEK), were not recovered in every simulation, as expected. However, edges with clear differences between the wild-type and mutant population were recovered well.

**Fig. 3:**
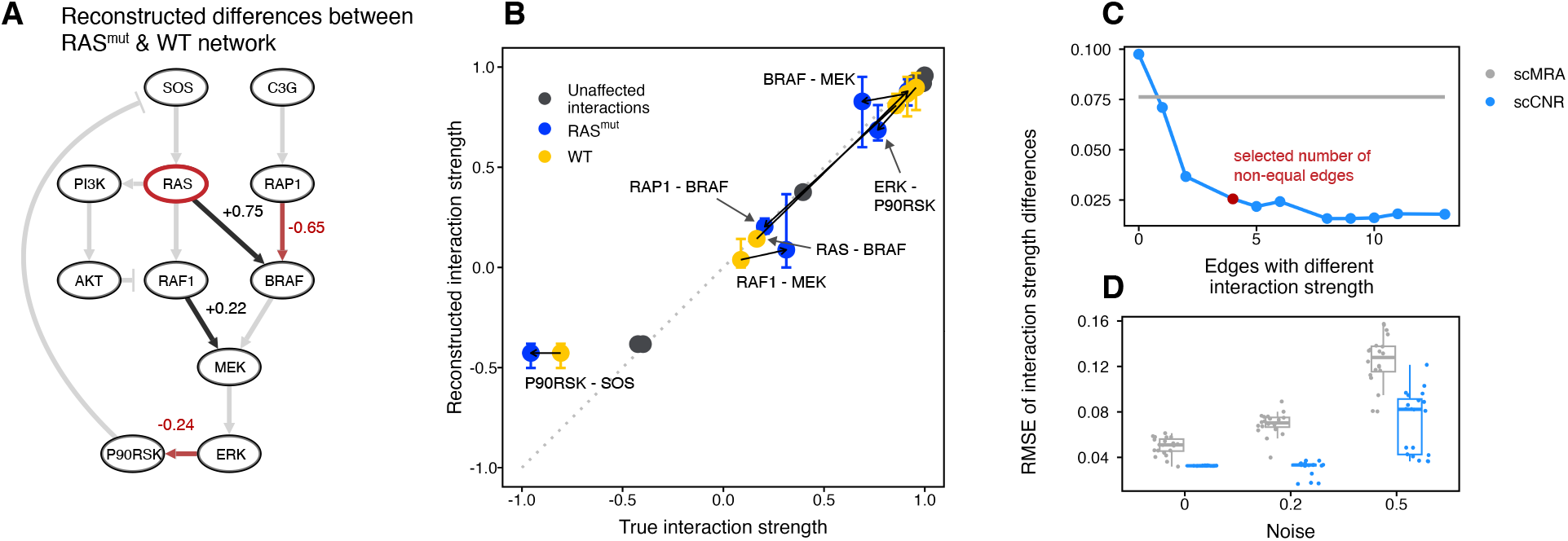
scCNR identifies signaling differences between cell populations. (**A**) Reconstructed differences between the RAS mutant and wild-type network. Gray background edges indicate the true network topology. Black and red edges highlight the positive and negative differences of the RAS mutant compared to the wild-type network. (**B**) Correlation between reconstructed and true interaction strengths averaged over 20 simulations. Blue points correspond to interaction strengths in the RAS mutant population, yellow in the wild-type population, and in gray points correspond to edges with the same interaction strength (difference below 0.1). Error bars visualize the range of reconstructed interaction strength values across simulations. Black lines connect corresponding wild-type and mutant interaction strengths. (**C**) RMSE of interaction strength differences between the two populations for increasing numbers of population-specific interaction strengths. (**D**) RMSE for joined scCNR (blue) with 4 population-specific edges or separate scMRA (gray) reconstructions of the network, with various levels of noise.

### 3.6. Joined network reconstruction with scCNR improves performance over reconstruction with scMRA

In addition to highlighting the major differences between populations, we hypothesized that scCNR might improve network reconstruction by pooling cells from all populations into a single optimization problem, hence increasing the power. To test this hypothesis we reconstructed the Orton network based on simulated wild-type and RAS mutant single-cell data of 500 cells in total, with 20% noise, and setting *η* to obtain networks with 13 edges. We quantified how well a combined reconstruction using scCNR recovers differences between mutant and wild-type networks, and compared this with two scMRA reconstructions where the wild-type and mutant populations are reconstructed independently. We calculated the difference of interaction strengths between the two populations (wild-type and mutant) for both the ground truth and the reconstructed networks and compared these. Fig. 3C shows the RMSE of the difference in interaction strengths as a function of the number of edges that have a different interaction strength between the two populations. The scCNR reconstructions (blue points) are consistently more accurate that the scMRA reconstructions (gray line), indicating the benefit of a joint network reconstruction. Increasing the number of differences above four results in little further improvement. With increasingly noisy input data, the increased performance of a joint reconstruction with scCNR becomes even more pronounced as seen in Fig. 3D and Fig. S3. Together, this shows that by pooling information from cells of multiple populations scCNR can more accurately reconstruct relevant differences between populations as compared to independent scMRA reconstructions.

### 3.7. scCNR detects signaling difference in response to EGFR inhibition

To further demonstrate the utility of scCNR we applied the method to real experimental single-cell measurements of EGFR-inhibitor or vehicle treated primary human keratinocytes, with the aim to quantify how prolonged EGFR inhibition affects signal transduction. While the EGFR pathway is well studied, how prolonged exposure to inhibitors alters signal transduction within the network is still an ongoing field of research (Klinger *et al*., 2013). In this experiment, we measured the abundance of 70 phosphorylated and total proteins of 282 cells using single-cell ID-seq (Eijl *et al*., 2018), involving 31 key nodes of the EGFR pathway, upon treatment with an EGFR inhibitor (AG1478) or vehicle (DMSO) control for 48 hours.

As expected, EGFR inhibition induced a reduction in phosphoprotein abundance of proteins downstream of EGFR, including RPS6 and AKT (Fig. S5A) (Fan *et al*., 2009; Phuchareon *et al*., 2015). We detect reduced levels of phospho-RB after treatment, while CDK4 and Cycline-E stayed active, which marks an arrest of cells in the early cell-cycle stage (Fig. S5A). Furthermore, EGFR inhibition pushes cells into a differentiated cell state, which is marked by a decrease in ITGB1. We also detect an increase of BMPR and it has been shown that BMP signaling goes up during differentiation (Eijl *et al*., 2018). EGFR inhibition additionally raises the level of phosphorylated p38, which has been linked to keratinocyte differentiation and cell-cycle arrest (Saha *et al*., 2014; Connelly *et al*., 2011; Adhikary *et al*., 2010; Efimova *et al*., 1998). Next to the increased phosphorylation of p38, rising levels of phosphorylated JNK have been reported for keratinocytes after EGFR-inhibitor treatment (Lu *et al*., 2011). Together, this demonstrates that the data contains biologically meaningful signal, and so we continued to model the underlying signaling differences between the two cell populations.

We applied scCNR to identify differences of the EGFR signaling pathway between the untreated and EGFR-inhibitor treated population. We ran scCNR with a prior-network topology of known protein interactions, and set the *θ*-parameter to identify twelve population-specific interaction strengths in the network (Fig. 4A) to balance model complexity and model fit (Fig. S5B).

**Fig. 4:**
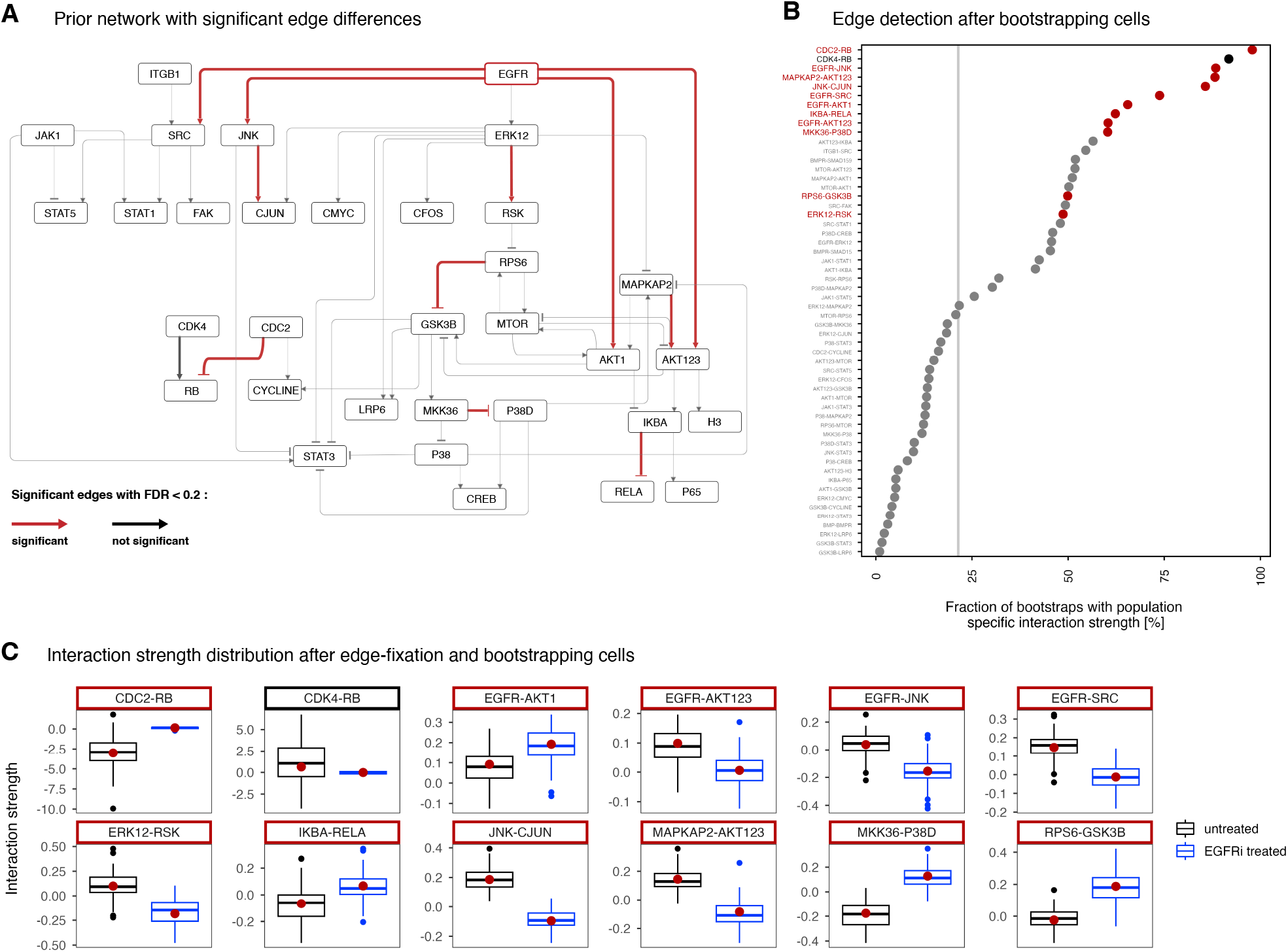
Analysis of signaling differences between untreated and EGFR-inhibitor treated keratinocytes. (**A**) Prior knowledge network highlighting reconstructed differences between EGFR-inhibitor treated and untreated cells. (**B**) Bootstrapping analysis to detect edges that are consistently identified to have population-specific strengths. After every bootstrap scCNR identified 12 population-specific interaction strengths and we counted their occurrence within all bootstrapped runs. Significant edges from A are coloured in red. The gray line corresponds to the theoretical expectation of randomly selecting an edge to have a population-specific interaction strength. (**C**) Distributions of interaction strengths from 1000 bootstrapping runs, for the untreated (black) and EGFR inhibitor treated cells (blue). The red dot indicates the recovered interaction strength from the whole data, as depicted in A.

To assess the statistical significance of the differences in interaction strengths between the treated and untreated populations identified by scCNR, we permuted the population labels and reconstructed the interaction strengths 1000 times, while fixing which of the interaction strengths can be population specific from the unpermuted solution. In this null-model we expect that the interaction strengths do not differ between populations. We calculated the empirical *p*-value as the fraction of permutations that led to a more extreme interaction strength difference than the one obtained from the unpermuted network reconstruction. At a 20% false positive rate, 11 of the 12 population-specific interaction strengths had significant differences between populations (Fig. 4A). As expected from an inhibition of EGFR, we predominantly recover differences directly linked to that node. Four of five outgoing edges from EGFR differ in interaction-strength between the cell populations. EGFR inhibition leads to cell-cycle arrest, and we recover signaling differences around the cell-cycle related node phospho-RB. However, phospho-RB follows a bimodal distribution in the untreated cell population and represents cells in different cell-cycle stages, which makes this observation hard to interpret. Nevertheless, the CDC2-RB edge is significant with a false discovery rate below 10% (Fig. S5C).

Next, we conducted a bootstrapping analysis to verify that the selection of edges is not primarily driven by outlier cells. We sub-sampled cells with replacement from both populations, repeated the network reconstruction 1000 times, and counted for each edge how often it was identified to be population specific. Edges around EGFR and phosho-RB consistently show up in the reconstruction, supporting the evidence of those differences between cell populations. Moreover, all differentially recovered edges are far above the theoretical occurrence of a randomly selected edge (Fig. 4B).

Finally, to investigate how reliable differences in interaction strength are *quantitatively*, we performed a bootstrapping analysis where we fixed all edges that were identified to differ in strength between populations, bootstrapped cells from both populations, and re-ran scCNR to recover interaction strengths. These 1000 bootstraps provided us with distributions of interaction strengths for both populations (Fig. 4C). Most interaction strengths showed clear differences between the cell populations. Notably, we observed that interactions with phospho-RB move to zero upon EGFR inhibition. Furthermore, among the four population-specific edges downstream of EGFR, three decrease in interaction strength following inhibition. Interactions with AKT123 and SRC diminish to zero and also edges further downstream of EGFR decrease in interaction strength upon EGFR inhibition, such as ERK12-RSK, JNK-CJUN or MAPKAP2-AKT123. From this, we conclude that despite the limitations imposed by the relatively small sample size, scCNR is able to detect significant and biologically meaningful differences in signal transduction between populations.

## 4 Discussion

How signal transduction differs between cell states, in disease, or upon treatment, plays a fundamental role in cell biology and has potential clinical implications. Reconstructing signaling networks has long been limited to bulk data, but recent advances in single-cell (phospho-)protein measurement technologies allows us to take advantage of many data points to study signaling networks on the single cell level. To this end, we developed single cell Modular Response Analysis (scMRA) and single cell Comparative Network Reconstruction (scCNR), which exploit the stochastic variability of protein abundances between cells as natural ‘perturbation experiments’. By penalizing network complexity and, in the case of scCNR, the differences in signaling between cell population, core signaling networks and their most relevant differences are found. Additionally, prior network information can easily be integrated. We extensively validated scMRA on simulated data and showed that the method works well even in presence of considerable noise and with as few as 500 cells. Additionally, we applied scCNR to a real-world single-cell ID-seq dataset of cells treated either with an EGFR inhibitor or with a vehicle control, and recovered biologically meaningful and expected signaling differences between the treated and untreated cell populations .

While our findings underscore the method’s capacity to identify relevant signaling differences, it is essential to also acknowledge the limitations. While scCNR explains phosphoprotein counts for every cell individually, it does rely on clustering of cells into discrete groups to infer population specific network parameters. In addition, some care needs to be taken in interpreting edges that the model proposes, as these might be indirect and mediated through unobserved nodes. Therefor, we expect the scCNR results to typically serve as a valuable foundation for generating hypotheses regarding mechanistic interactions, which can subsequently be subjected to further in-depth exploration. Future progress in single-cell protein measurement techniques will enhance the detection of cell-to-cell variability and will improve network inference with scCNR.

In contrast to the classical MRA and CNR formulation, the scMRA and scCNR optimization problems are typically over-determined since there are many more linear equations - one for every cell and every protein - than possible edges. Nonetheless, the computational complexity of the optimization problem increases exponentially with the number of nodes in the network. However, biological networks are often sparsely connected, and sparse solutions can generally be found efficiently. For instance, the Orton model can be solved in under 20 seconds for 500 cells with 20% noise and 13 edges on a standard laptop. The number of nodes in the network and the number of edges and population-specific interaction strengths influence the search space and therefore the run time. In cases where many interactions have to be inferred the run time can be reduced drastically by incorporating prior network information or by setting hyperparameter settings that limit the number of model parameters that need to be inferred.

Several methods to reconstruct population specific networks from single cell data have been proposed before(Rukhlenko *et al*., 2022; Brandt *et al*., 2019; Kumar *et al*., 2020). However, these methods use the single cell nature of the data only to cluster cells in distinct groups, and then aggregate the cells within one population. As such, they do not exploit the within population variation in signaling activity and they require many perturbations for network inference. DREMI introduced the concept of using cell-to-cell variation in protein activity to gain insight into the strength of protein interactions (Krishnaswamy *et al*., 2014), but this method only considers protein pairs, and does not reconstruct networks. Compared to these methods, the advantages of scCNR are that it uses variability within the single-cell population, with the possibility to easily integrate perturbations, prior networks and various cell populations, to identify cell population specific signaling networks.

In conclusion, scMRA enables the reconstruction of signal transduction networks from single-cell data, and scCNR additionally recovers the most relevant differences in signaling between cell populations. We envision that scCNR will help in the recovery of signaling networks in diverse biological settings, especially by shedding light on clinically relevant signaling differences, e.g., between cell states and upon treatment.

## Notes

### Competing Interest Statement

The authors have declared no competing interest.

