## Supplementary material for "Reconstructing and comparing signal transduction networks from single cell protein quantification data"

### 5 Supplementary Material

#### 5.1. Supplementary Methods

##### 5.1.1. Evaluating scMRA and scCNR

For the analyses presented in Fig. 2 E and Fig. 3, to compare networks with the same number of edges, we reconstructed networks iteratively and varied the $\eta$ parameter until we reached a network of 13 edges. 13 edges also show a major reduction in the mean squared error of the objective function while keeping a sparse network topology (Fig. S1B). For the evaluation of scCNR in Fig. 2 and Fig. S4 we followed the same procedure and compared only reconstructed networks with 13 edges. Balancing the model complexity and the reduction in the mean squared error resulted in 4 population-specific interaction strengths for the scCNR analysis of the RAS mutant population as well as for the BRAF mutant population (Fig. S2). To evaluate how well scCNR detects differences we calculated the RMSE of the interactions strenghts differences in Fig. 3C,D. scCNR results in population-specific interaction strenghts that share the same network topology. We calculated the difference for all edges in the network between the two populations. We did the same for the ground truth data of the wild type and mutant simulations. To compare the true with the recovered differences we calculated the RMSE between these two sets. The increased performance of scCNR to identify signaling differences is exemplified in network reconstructions based on very noisy (50%) input data (Fig. S3). We performed reconstructions based on 20 simulation runs, and calculated the average interaction strength differences of these 20 reconstructions. The two main interaction differences for the comparison of the RAS mutant and wild-type network, RAS-BRAF and RAP1-BRAF, are discovered in both the scCNR and scMRA approach. For scCNR we find two solutions across the 20 simulations (bimodal RMSE of interaction strength differences (Fig. 3D))). One solution models the network correctly with edges from RAS and BRAF, while the other solution identifies wrong interactions from SOS, C3G. However, scMRA detects many false positive interaction differences, such as RAS-SOS (present in all 20 simulations), RAP1-C3G (present in 70% of the simulations) and P90RSK-SOS (present in 90%) with a wrong sign of the interaction strength difference.

##### 5.1.2. Cell culture and reagents

Primary pooled human epidermal stem cells derived from foreskin were obtained from Lonza. Cells were cultured and expanded as previously reported (Gandarillas and Watt, 1997). Briefly, cells were cultured on a feeder layer of J2-3T3 cells in FAD medium (Ham's F12 medium/Dulbecco's modified Eagle medium (DMEM) (1:3) supplemented with 10% batch tested fetal calf serum (FCS) and a cocktail of 0.5 µg/ml of hydrocortisone, 5 µg/ml of insulin, 0.1 nM cholera enterotoxin, and 10 ng/ml of epidermal growth factor) supplemented with Rock inhibitor (Y-27632, 10 µM). J2-3T3 cells were cultured in DMEM containing 10% bovine serum and inactivated with Mitomycin C (SCBT) upon seeding the epidermal stem cells. For experiments epidermal stem cells were transferred to Keratinocyte Serum Free Medium (KSFM) supplemented with 0.2 ng/ml Epidermal Growth Factor and 30 µg/ml bovine pituitary extract from Life Technology until 70% confluent. Cells were treated with AG1478 (10 µM, Calbiochem) or vehicle (DMSO) for 48 hours prior to harvesting and fixation. All media were supplemented with 1% penicillin/streptomycin antibiotics.

##### 5.1.3. Antibody conjugation with dsDNA barcodes

Antibodies and dsDNA were functionalized and conjugated as described (Buggenum *et al.*, 2016). The antibody panel was used as in (Eijl *et al.*, 2018) and details are provided in supplemental Table 1 of this paper. In short, antibodies were functionalized with NHS-s-s-PEG4-tetrazine (Jena Bioscience) in a ratio of 1:10 in 50 mM borate buffered Saline pH 8.4 (150 mM NaCl). Then, N3-dsDNA was produced and functionalized with DBCO-PEG12-TCO (Jena Bioscience) in a ratio of 1:25 (oligo list). Finally, purified functionalized antibodies were conjugated to purified functionalized DNA by 4-hour incubation at room temperature in borate buffered saline pH 8.4 in a ratio of 4:1 respectively. The reaction was quenched with an excess of 3,6-diphenyl tetrazine. The conjugation efficiency and quality were checked on an agarose gel, confirming that a substantial amount of DNA conjugated with the antibody. Ultimately, conjugates were equally pooled for staining's in scID-seq.

##### 5.1.4. Immunostaining and single-cell sorting

The scID-seq workflow was performed essentially as described in (Eijl *et al.*, 2018), In brief, cells ($> 3x10^6$) were harvested with trypsin and cross-linked in suspension by incubating for 10 minutes with 4% paraformaldehyde (PFA) in PBS following a quenching step of 5 minutes with 125 mM Glycine in PBS. Removal of PFA and Glycine occurred through washing twice with wash buffer (0.1x Pierce™ Protein-Free Blocking Buffer from Thermo in PBS). Then, cells were blocked in 500µl blocking buffer (0.5x 0.1x Pierce™ Protein-Free Blocking Buffer, 200 µg/ml boiled salmon sperm DNA, 0.1% Triton-X 100, in PBS) at room temperature for 30-60 min. Staining with the conjugate mix occurred overnight at 4°C in 500 µl blocking buffer and washed 3x in 5ml wash buffer. Individual cells were sorted with the BD FACSAria SORP flow cytometer (BD biosciences) in 96 well PCR plates containing 1µl release buffer (10 mM DTT in 15mM Tris, pH 8.8) and 7µl Vapor-lock (Qiagen). Plates were stored at -20°C until use.

##### 5.1.5. Barcoding and library preparation for next generation sequencing

For the library preparation 3 PCR steps were performed to amplify the antibody barcodes and to add barcodes specific for the well and the plate of each cell. The barcoding occurred with the same sequences used in ID-seq (Buggenum *et al.*, 2016). For the first PCR step 15 cycles were run after adding to each well a 4 µl reaction mix containing the Herculase II Fusion DNA Polymerase (Agilent), dNTPs, 5x Herculase buffer and 0.1µM amplification primers (Forward `5'-CACGACGCTCTTCCGATCT-3'`, Reverse `5'-TCGCTTATCTGTTGACTGAT-3'`). Directly after the first PCR step, 5 extra cycles were run after adding 1 µl mix containing Herculase buffer 0.2 µM forward amplification primer and 0.2 µM reverse well barcoding primer. Then all material was pooled per plate, Vapor-lock was removed and a clean-up was performed with the QIAquick PCR Purification Kit, an EXO1 treatment to degrade remaining primers followed by another purification. Another 5 cycles were run in PCR 3 with a 20 µl reaction containing pooled and purified plate sample and 0.1 µM plate barcoding primers (Fw_long `5'-AATGATACGGCGACCACCGAGATCTACACTCTTTCCCTACACGACGCTCTTCCGATCT-3'` and specific plate reverse). After repeating the clean-up, the libraries were checked on agarose gel and with the Bioanalyzer (Agilent) to confirm the size of the DNA fragments (expected size around 185 bp).

##### 5.1.6. scID-seq data analysis

Sequence data from the NextSeq500 (Illumina) was demultiplexed using bcl2fastq software (Illumina). Then, all reads were processed using our dedicated R-package (van Buggenum et al., 2018). In short, the sequencing reads were split using a common "anchor sequence" identifying the position of the UMI sequence, Barcode 1 (antibody specific) and Barcode 2 (well specific) sequence. After removing all duplicate reads, the number of UMI sequences were counted per barcode 1 and 2. Finally, barcode 1 ("antibody)" and barcode 2 ('well') sequences were matched to the corresponding. For scID-seq, a threshold was set based on the total UMI count per well difference between high- and low-quality cells or empty wells. Antibodies with a median of $< 10$ counts per cell were removed from the dataset. The data was normalized using TMM with the default A-values (Robinson and Oshlack, 2010). Since the dataset contained a defined panel of measured proteins we reduced the M-values to 5% to keep a sufficient set of proteins for the normalization. Differential abundance of phosphoproteins between the untreated and EGFR inhibitor treated cell population was calculated with DESeq2.

The signalling differences between the untreated and EGFR-inhibitor treated cell populations were reconstructed using scCNR. The data displayed considerable heterogeneity, especially the cell cycle marker phospho-RB had a bimodal distribution separating cells into an actively cycling and non-cycling subgroup for the untreated population. Since the number of cells was limited (164 untreated and 118 treated cells) we nevertheless did not further divide the untreated group into sub-populations. Outliers were removed separately for each population by removing cells with a protein abundance difference from the population mean that exceeded three standard deviations. This reduced the total cell counts to 117 untreated and 95 treated cells and resulted in recovered network differences that appeared stable towards outliers (Fig. 4). The EGFR network topology is well studied and we started the analysis with a prior network. We added canonical interaction as well as known interactions from PhoshoSitePlus® (2022-10-21) to the network. We then estimated the penalty for the number of non-equal edges ($\theta$) to reconstruct a network with 12 different edges. The permutation analysis to assess the statistical significance of interaction strength differences depicted in (Fig. 4A) was performed by randomly shuffling cell state labels. The network topology and the interactions that differ were fixed by constraining all indicator variables to be the value they had in the original model. Subsequently, scCNR was re-run 1000 times to obtain the interaction strengths for the permuted data. This way we obtain a null-distribution of interaction-strength-deviations from the population mean (Fig. S5C). The bootstrap analysis to assess the robustness of interaction-strength-difference detection in Fig. 4B was performed by randomly sampling 212 cells (i.e. the original number of cells) with replacement, and rerunning scCNR with its original parameters. This procedure was repeated 1000 times. From this, the fraction of bootstraps in which an edge is predicted to be different can be obtained. In Fig. 4C we fixed the interaction strengths differences found in Fig. 4A by enforcing the corresponding indicator constraints in the scCNR model. We then performed the bootstrapping analysis similarly as above by sampling 212 cells with replacement and repeating the bootstrapping 1000 times. From this we obtained distributions of interaction strengths in both populations for the 12 population-specific interaction strengths.

#### 5.2. Derivation of the single-cell MRA equation

We assume we have a dataset of $N_c$ cells, in which we have measured the activity $x_i$ and total abundance $x_i^{\text{tot}}$ of $N_n$ proteins, which are the nodes of the signaling network we want to reconstruction. Optionally, a set of $N_p$ perturbations by e.g. small molecule inhibitors was performed.

The activity of node $i$ in cell $a$, $x_{i,a}$ can be described as the population average activity of node $i$, $\langle x_i \rangle$ plus some deviation $\Delta_a x_i$.

$$x_{i,a} = \langle x_i \rangle + \Delta_a x_i \tag{2}$$

and similarly for the total amount of protein $x_i^{\text{tot}} = \langle x_i^{\text{tot}} \rangle + \Delta_a x_i^{\text{tot}}$. We assume that each cell individually is in (quasi) steady state, which implies that

$$\frac{\mathrm{d}x_{i,a}}{\mathrm{d}t} = f_i\left(\mathbf{x}_a, x_{i,a}^{\text{tot}}, \mathbf{p}\right) \equiv f_{i,a} \approx 0 \quad \forall a, i. \tag{3}$$

Similarly, the assumption that the bulk average is a steady state implies that

$$\frac{\mathrm{d}\langle x_i \rangle}{\mathrm{d}t} = f_i\left(\langle \mathbf{x} \rangle, \langle x_i^{\text{tot}} \rangle, \mathbf{p}\right) \equiv \langle f_i \rangle = 0 \quad \forall i. \tag{4}$$

Perturbations to the network, such treatment with small molecule inhibitors, can be modelled as a perturbation to the parameters in the unperturbed reference condition $\mathbf{p_0}$, i.e. $\mathbf{p} = \mathbf{p}_0 + \Delta\mathbf{p}$

The first order Taylor expansion of $f_{i,a}$ around $\langle x_i \rangle$ is given by:

$$f_{i,a} \approx \langle f_i \rangle + \sum_{j=1}^{N_n} \frac{\partial f_i}{\partial x_j} \cdot \Delta_a x_j + \frac{\partial f_i}{\partial x_i^{\text{tot}}} \cdot \Delta_a x_i^{\text{tot}} + \sum_{m=1}^{N_p} \frac{\partial f_i}{\partial p_m} \cdot \Delta p_m, \tag{5}$$

where we made the assumption that $\frac{\partial f_i}{\partial x_j^{\text{tot}}} = 0$ if $i \neq j$. This implies that the activity of a protein is not *directly* affected by changes in the total abundance of other proteins, which in many cases is a biologically sound assumption. Also note that $\frac{\partial f_i}{\partial p_m} \neq 0$ only for those perturbations $p_m$ that *directly* affect node $x_i$, such as a targeted inhibitor and its kinase target. Dividing everything by $\langle x_i \rangle \frac{\partial f_i}{\partial x_i}$ and using that $f_{i,a}$ and $\langle f_i \rangle$ are 0 this gives

$$\begin{aligned} 0 \approx &\frac{\Delta_a x_i}{\langle x_i \rangle} + \sum_{j \neq i} \frac{\langle x_j \rangle}{\langle x_i \rangle} \left( \frac{\partial f_i}{\partial x_j} \Big/ \frac{\partial f_i}{\partial x_i} \right) \cdot \frac{\Delta_a x_j}{\langle x_j \rangle} + \\ &\frac{\langle x_i^{\text{tot}} \rangle}{\langle x_i \rangle} \left( \frac{\partial f_i}{\partial x_i^{\text{tot}}} \Big/ \frac{\partial f_i}{\partial x_i} \right) \cdot \frac{\Delta_a x_i^{\text{tot}}}{\langle x_i^{\text{tot}} \rangle} + \\ &\frac{p_m}{\langle x_i \rangle} \left( \frac{\partial f_i}{\partial p_m} \Big/ \frac{\partial f_i}{\partial x_i} \right) \cdot \frac{\Delta p_m}{p_m}. \end{aligned} \tag{6}$$

the implicit function theorem states that

$$\frac{\partial f_i}{\partial x_j} \Big/ \frac{\partial f_i}{\partial x_i} = -\frac{\partial x_i}{\partial x_j}, \quad \frac{\partial f_i}{\partial x_i^{\text{tot}}} \Big/ \frac{\partial fi}{\partial xi} = -\frac{\partial x_i}{\partial x_i^{\text{tot}}}, \text{ and } \frac{\partial f_i}{\partial p_m} \Big/ \frac{\partial f_i}{\partial x_i} = -\frac{\partial x_i}{\partial p_m},$$

so that we end up with the equation:

$$R_{i,a} \approx \sum_{j \neq i} r_{ij} \cdot R_{j,a} + s_i \cdot R_{i,a}^{\text{tot}} + \sum_m p_{im} \quad \forall a, i. \tag{7}$$

where:

$$r_{ij} \equiv \frac{\langle x_j \rangle}{\langle x_i \rangle} \frac{\partial x_i}{\partial x_j} \tag{8}$$

is the interaction strength between node $j$ and node $i$, with $r_{ii} \equiv -1$.

$$s_i \equiv \frac{\langle x_i^{\text{tot}} \rangle}{\langle x_i \rangle} \frac{\partial x_i}{\partial x_i^{\text{tot}}}, \tag{9}$$

is the sensitivity of $x_i$ to changes in it's total abundance,

$$p_{im} \equiv \frac{p_m}{\langle x_i \rangle} \frac{\partial x_i}{\partial p_m} \frac{\Delta p_m}{p_m} \tag{10}$$

is the direct effect that perturbation $m$ has on the activity of node $i$,

$$R_{i,a} \equiv \frac{\Delta_a x_i}{\langle x_i \rangle} \tag{11}$$

is the deviation of the activity of node $i$ from the population average and

$$R_{i,a}^{\text{tot}} \equiv \frac{\Delta_a x_i^{\text{tot}}}{\langle x_i^{\text{tot}} \rangle} \tag{12}$$

is the deviation of the total abundance of protein $i$ from the population average.

In matrix-form, equation 7 reads:

$$\mathbf{r} \cdot \mathbf{R} + \mathbf{s} \cdot \mathbf{R}^{\text{tot}} + \mathbf{p} \approx 0 \tag{13}$$

$r_{ij}$, $s_i$ and $p_{i,m}$ are the unknown quantities that we want to estimate based on the measurements $R_{i,a}$ and $R_{i,a}^{\text{tot}}$. For this, we have system of $N_n \times N_c$ equations (one for each cell for each node). In theory, there are $N_n \times N_n + N_n + N_n \times N_p$ unknowns. However, in practice we will know for the vast majority of the $N_n \times N_p$ elements of $p_{im}$ that they are 0, because most nodes are not affected by most inhibitors.

#### 5.3. MIQP formulation of single-cell MRA

Our aim is to find a network model that accurately describes the data, without being overly complex. To this end, similar as for "bulk" Comparative Network Reconstruction (Bosdriesz *et al.*, 2018), we formulate the problem as a Mixed Integer Quadratic Programming (MIQP) problem. We aim to solve Equation 7, while placing an $L_0$ penalty on the presence of an edge. Given $N_n$ nodes in the network,

$N_c$ cells measured and for $i,j \in \{1,2,\ldots,N_n\}$, $n \in \{1,2,\ldots,N_p\}$ and $a \in \{1,2,\ldots,N_c\}$, the MIQP problem then reads as follows:

$$\begin{aligned}
\text{minimize} : & \sum_{a,i,j} \left[(1-\eta)\cdot \epsilon_{i,a}^2 + \eta \cdot I_{ij}^{\text{edge}}\right] \\
\text{subject to} : & \sum_{j=1}^{N_n} r_{ij}\cdot R_{j,a} + s_i \cdot R_{i,a}^{\text{tot}} + \sum_{m=1}^{N_p} p_{im} = \epsilon_{i,a} && \forall a,i \\
& r_{ii} = -1 \\
& s_i > 0 && \forall i \\
& I_{ij}^{\text{edge}} = 0 \Rightarrow r_{ij} = 0 && \forall i,j \\
& I_{ij}^{\text{edge}} \in \{0,1\}
\end{aligned}$$

where $I_{ij}^{\text{edge}}$ are so-called indicator constraints indicating the presence or absence of an edge from node $j$ to node $i$, which are used to implement the $L_0$ penalty. The hyperparameter $\eta$ is used to tune the complexity of the model. We also make the assumption that an increase in total protein abundance increases node activity: $s_i > 0$.

In some cases, for some nodes only the activity but not their total abundance is measured, i.e. $R_{i,a}^{\text{tot}}$ is not measured for protein $i$. In that situation we cannot estimate $s_i$. The most straightforward solution is to ignore the $s_i \cdot R_{i,a}^{\text{tot}}$ term. As a results, the residuals for these nodes will be higher, on average. To account for this, we can weight the residuals of "complete" and "incompletely" nodes differently. Specifically, this requires finding a factor $\alpha$ (between 0 and 1) such that $\alpha \cdot \langle \epsilon_{\text{i}}^2 \rangle = \langle \epsilon_c^2 \rangle$ with the subscript referring to incomplete and complete nodes, respectively.

This requires estimating the contribution of the $s_i \cdot R_{i,a}^{\text{tot}}$-terms to explaining the response, which is not possible exactly. However, we can get an upper and lower bound:

- For the upper bound, we assume there is no contribution from local responses $r_{ij}$. We can obtain this by calculating the residuals with $r_{ij} = 0$. This basically means regressing $R_{i,a}$ against $R_{i,a}^{\text{tot}}$.
- For the lower bound, we assume there is a maximal contribution from local responses $r_{ij}$. We obtain this by not penalizing edges at all in the reconstruction, i.e. by setting $\eta = 0$.

We then select $\alpha$ by taking the mean of the upper and the lower bound.

#### 5.4. MIQP formulation of single-cell CNR

In many instances, we are interested in comparing the signaling networks between $N_{\text{cs}}$ different cell populations $\xi$, with each cell state containing $N_c^\xi$ $\xi \in \{1,\ldots,N_{cs}\}$ cells. Similar as for bulk Comparative Network Reconstruction (Bosdriesz *et al.*, 2018), the main idea is to formulate the MRA equation 7 on a per-cell population basis, and solve these simultaneously in an MIQP, while placing an $L_0$-penalty on quantitative differences between the networks. Note that in such a case, the deviations of total and phosphoprotein abundances should be calculated relative to the average of the cell population $\xi$, i.e. $x_{i,a} = \langle x_i^\xi \rangle + \Delta_a x_i$.

To that end, we can formulate Equation 5.3 on a per-cell state basis.

$$R_{i,a} \approx \sum_{j \neq i} r_{ij}^\xi \cdot R_{j,a} + s_i^\xi \cdot R_{i,a}^{\text{tot},x} + p_{im}^\xi \quad \forall a,i,x. \tag{14}$$

where the $\xi$-superscript refers to the cell population. The variables than become:

$$r_{ij}^\xi \equiv \frac{\langle x_j^\xi \rangle}{\langle x_i^\xi \rangle} \frac{\partial x_i^\xi}{\partial x_j^\xi} \tag{15}$$

$$s_i^\xi \equiv \frac{\langle x_i^{\text{tot},\xi} \rangle}{\langle x_i^\xi \rangle} \frac{\partial x_i}{\partial x_i^{\text{tot}}} \tag{16}$$

$$R_{i,a} \equiv \frac{\Delta_a x_i}{\langle x_i^\xi \rangle} \tag{17}$$

$$R_{i,a}^{\text{tot}} \equiv \frac{\Delta_a x_i^{\text{tot}}}{\langle x_i^{\text{tot},\xi} \rangle} \tag{18}$$

The full scCNR MIQP formulation becomes:

$$\begin{aligned}
\text{minimize} : & \sum_{a,i,j,m} \left[(1-\eta)\cdot \epsilon_{i,a}^2 + \eta \cdot I_{ij}^{\text{edge}} + \theta(I_{ij}^r + I_i^s + I_{im}^p\right] && \forall a,i,j. \\
\text{subject to} : & \sum_{j=1}^{N_n} r_{ij}^\xi \cdot R_{j,a} + s_i^\xi \cdot R_{i,a}^{\text{tot}} + \sum_{m=1}^{N_p} p_{im}^\xi = \epsilon_{i,a} && \forall a,i \\
& r_{ii}^\xi = -1 \\
& s_i^\xi > 0 && \forall i \\
& I_{ij}^{\text{edge}} = 0 \Rightarrow r_{ij} = 0 && \forall i,j \\
& I_{ij}^r = 0 \Rightarrow r_{ij}^\xi = \langle r_{ij} \rangle && \forall i,j \\
& I_i^s = 0 \Rightarrow s_i^\xi = \langle s_i \rangle && \forall i \\
& I_{im}^p = 0 \Rightarrow p_{im}^\xi = \langle p_{im} \rangle && \forall i,m \\
& I_{ij}^{\text{edge}}, I_{ij}^r, I_i^s, I_{im}^p \in \{0,1\}
\end{aligned}$$

where

$$\begin{aligned}
\langle r_{ij} \rangle &\equiv \frac{1}{N_{cs}} \sum_\xi r_{ij}^\xi, \\
\langle s_i \rangle &\equiv \frac{1}{N_{cs}} \sum_\xi s_i^\xi, \text{ and} \\
\langle p_{im} \rangle &\equiv \frac{1}{N_{cs}} \sum_\xi p_{im}^\xi
\end{aligned} \tag{19}$$

5.5. Supplementary Figures

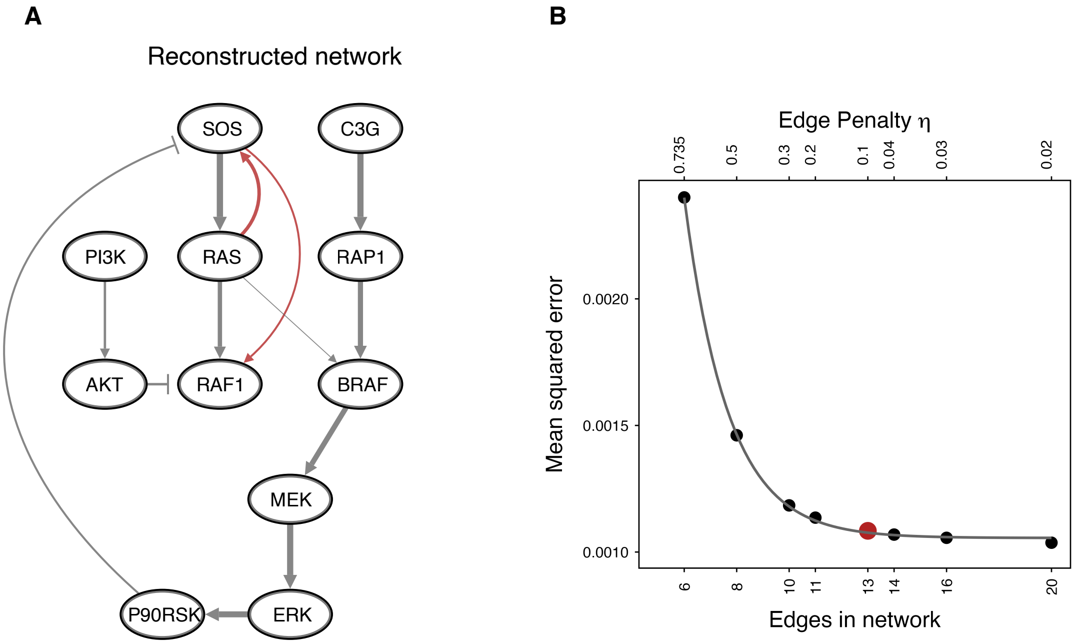


Fig. S1: **Exemplary scMRA network reconstruction with 13 edges**. (**A**) Reconstructed network for a simulation of the Orton Model as in (Fig. 2C) with 1000 cells and 20% noise. (**B**) To balance a good network reconstruction and a simple network topology (focussing on the main edges) we recommend to pick an $\eta$ at the "elbow" of the curve for the final reconstruction, where the fitting error of the model shows minimal improvement with increasing network complexity. The network of the Orton Model contains 13 edges, which is marked in red.

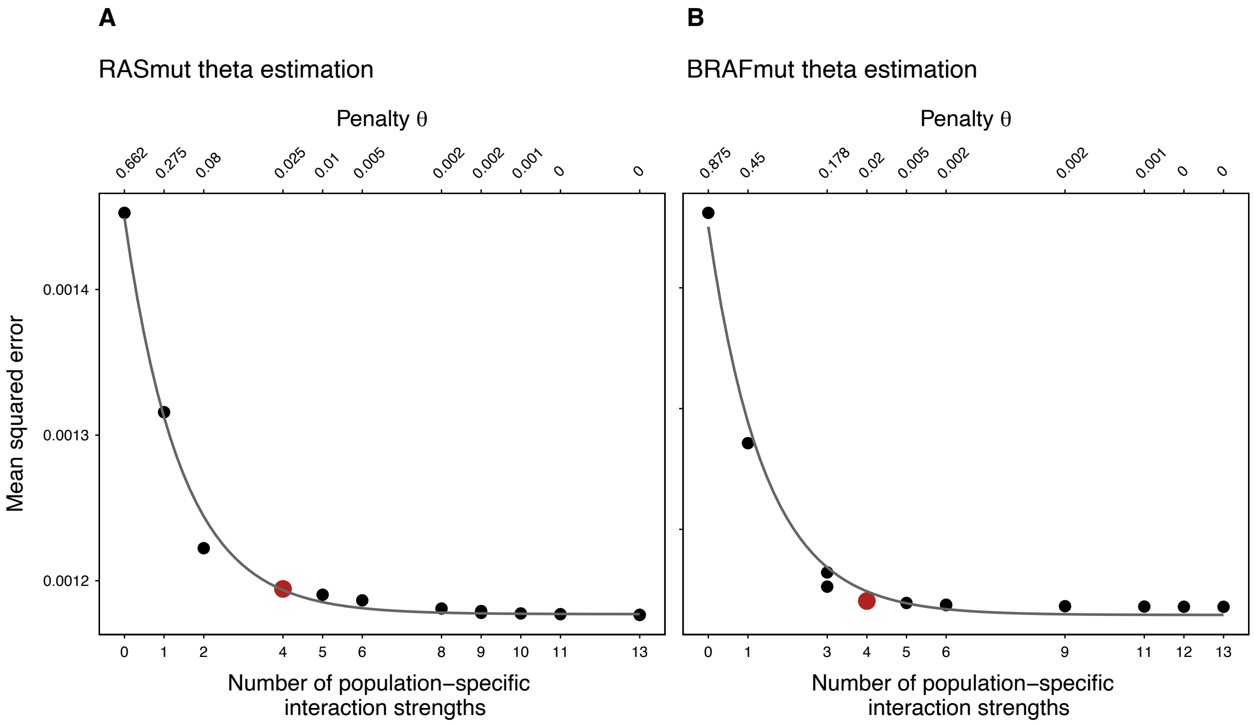


Fig. S2: **Estimation of the penalty on number of different edges ($\theta$).** For the reconstruction of the RAS mutant network (**A**) and the BRAF mutant network (**B**) we chose the parameter to reconstruct 4 edges with population-specific interaction strengths.

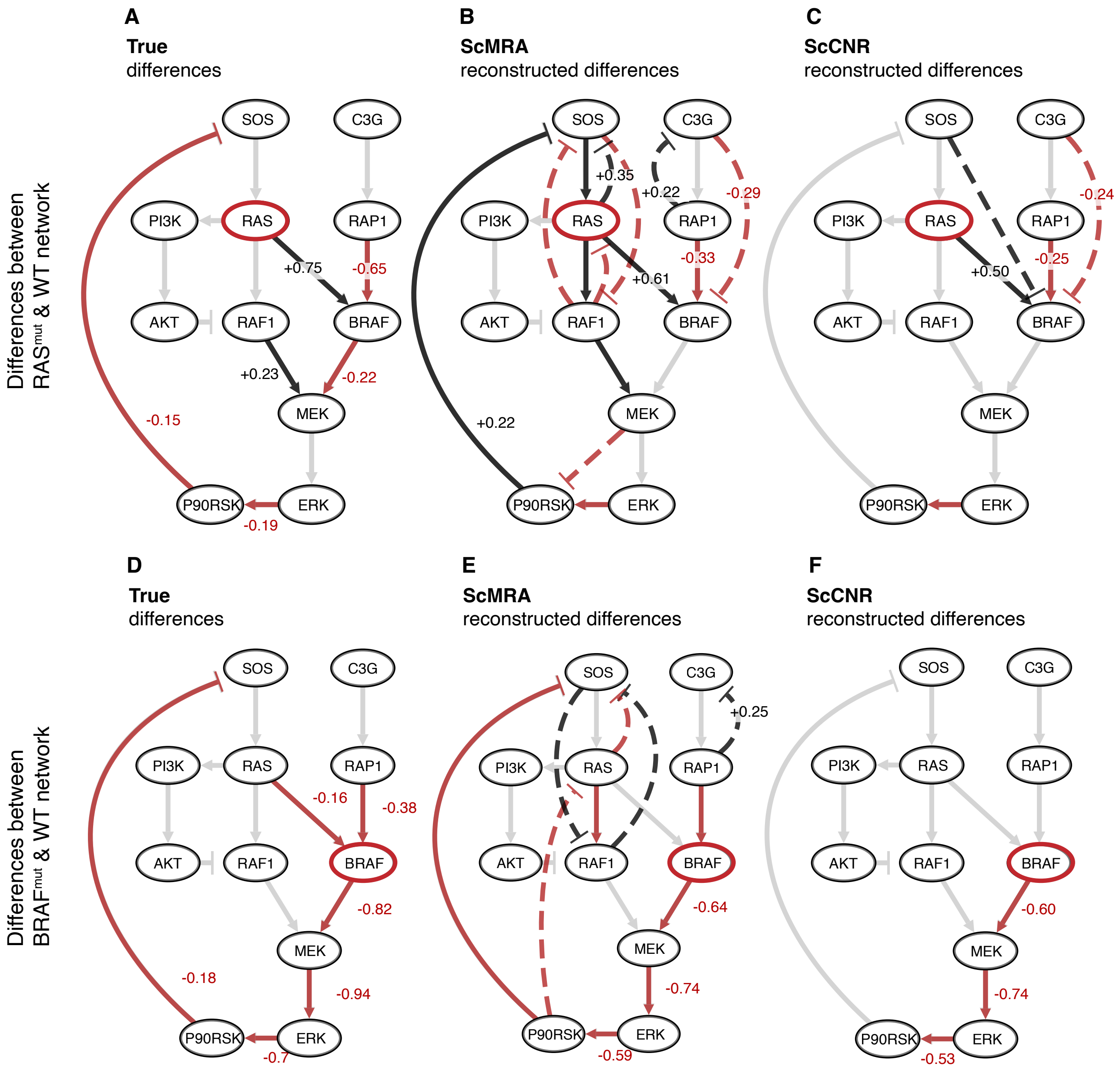


Fig. S3: **scCNR network reconstruction with high noise (50%).** The first row shows network differences between the Ras mutant and wild-type population, the second row between the BRAF mutant and wild-type population. From left to right population-specific interactions in the network for the Orton model (**A**,**D**), combined scCNR reconstruction (**B**,**E**), and two separate scMRA reconstructions (**C**,**F**). Every network was reconstructed from 20 different simulations and interaction strengths were averaged. Interactions that are stronger in the mutant compared to the wild type are in black and edges with an decreased strength are in red. Edges with a strength-difference above 0.1 are visualized and differences above 0.2 are annotated. Dashed lines represent false positive edges. The true network topology is depicted in gray in the background.

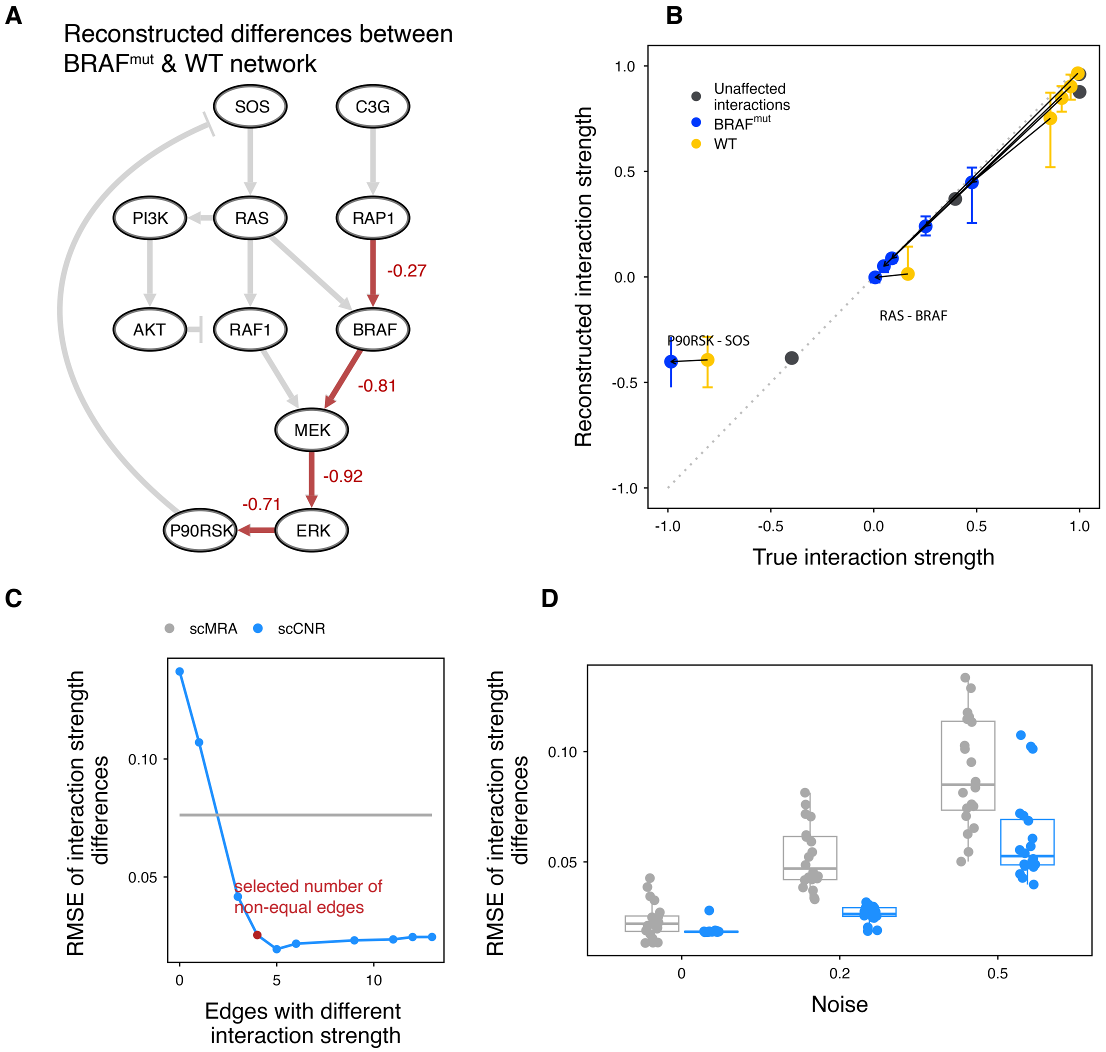


Fig. S4: **Reconstructing signalling differences between the BRAF mutant and wild-type cell population.** (**A**) Reconstructed differences between the BRAF mutant and wild-type network from two exemplary cell populations of 250 cells with 20% noise. In grey in the background is the true network topology. In black/ red positive and negative differences of the BRAF mutant network compared to the wild-type network are highlighted. (**B**) Correlation of reconstructed (average of 20 simulations with 250 cells per population and 20% noise) versus true interaction strength for edges in the BRAF mutant (blue), wild-type (yellow) population and in grey edges with the same interaction strength in both populations (difference below 0.1). Error bars on every point visualize maximum variation of the reconstructed interaction strengths. Black lines connect the wild type and mutant interaction strengths. (**C & D**) Reconstruction of differences between wild type and mutant cell population measured by RMSE of interaction strength differences. (C) Effect of modelling increasing number of non-equal interaction strengths on the RMSE of interaction strenght differences. (D) RMSE of differences for joined/ separate reconstructions of the network with various levels of noise.

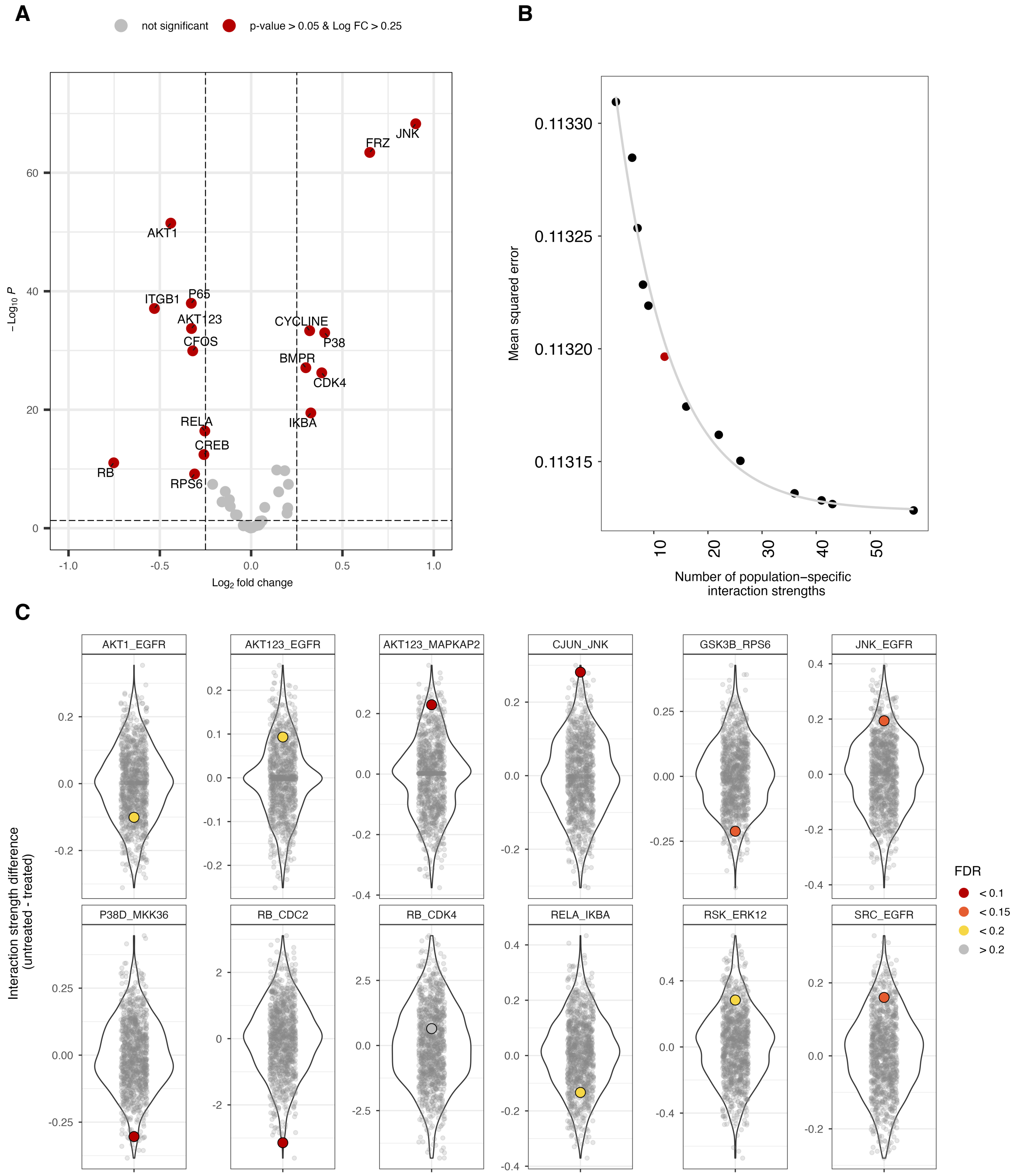


Fig. S5: **Analysis of signaling differences between untreated and EGFR-inhibitor treated keratinocytes.** (**A**) Volcano plot of proteins from the ID-seq panel. Proteins with a corrected p-value below 0.5 and log fold change higher than 0.3 are annotated. (**B**) Mean squared error of the scCNR analysis for an increasing number of population-specific interaction strengths. (**C**) Permutation analysis to estimate significance of detected edge differences. We randomly permuted cell population labels (treated/ untreated), fixed edges that were found to be different in the original data and repeated scCNR 1000 times. Colour coded dots represent the origionally recovered interaction strength differences and their false discovery rate.
